# Genome-wide identification of long non-coding RNAs and their regulatory networks involved in *Apis mellifera ligustica* response to *Nosema ceranae* infection

**DOI:** 10.1101/643627

**Authors:** Rui Guo, Huazhi Chen, Yu Du, Dingding Zhou, Sihai Geng, Haipeng Wang, Zhiwei Zhu, Caiyun Shi, Jieqi Wan, Cuiling Xiong, Yanzhen Zheng, Dafu Chen

**Author notes:** These authors contributed equally to this work (Rui Guo, Huazhi Chen).

## Abstract

Long non-coding RNAs (lncRNAs) are a diverse class of transcripts that structurally resemble mRNAs but do not encode proteins, and lncRNAs have been proved to play pivotal roles in a wide range of biological processes in animals and plants. However, knowledge of expression pattern and potential role of honeybee lncRNAs response to *Nosema ceranae* infection is completely unknown. Here, we performed whole transcriptome strand-specific RNA sequencing of normal midguts of *Apis mellifera ligustica* workers (Am7CK, Am10CK) and *N. ceranae*-inoculated midguts (Am7T, Am10T), followed by comprehensive analyses using bioinformatic and molecular approaches. A total of 6353 *A. m. ligustica* lncRNAs were identified, including 4749 conserved lncRNAs and 1604 novel lncRNAs. These lncRNAs had low sequence similarities with other known lncRNAs in other species; however, their structural features were similar with counterparts in mammals and plants, including shorter exon and intron length, lower exon number, and lower expression level, compared with protein-coding transcripts. Further, 111 and 146 *N. ceranae*-responsive lncRNAs were identified from midguts at 7 day post inoculation (dpi) and 10 dpi compared with control midguts. 12 differentially expressed lncRNAs (DElncRNAs) were shared by Am7CK vs Am7T and Am10CK vs Am10T comparison groups, while the numbers of unique ones were 99 and 134, respectively. Functional annotation and pathway analysis showed the DElncRNAs may regulate the expression of neighboring genes by acting in *cis*. Moreover, we discovered 27 lncRNAs harboring eight known miRNA precursors and 513 lncRNAs harboring 2257 novel miRNA precursors. Additionally, hundreds of DElncRNAs and their target miRNAs were found to form complex competitive endogenous RNA (ceRNA) networks, suggesting these DElncRNAs may act as miRNA sponges. Furthermore, DElncRNA-miRNA-mRNA networks were constructed and investigated, the result demonstrated that part of DElncRNAs were likely to participate in regulating the material and energy metabolism as well as cellular and humoral immune during host responses to *N. ceranae* invasion. Finally, the expression pattern of 10 DElncRNAs was validated using RT-qPCR, confirming the reliability of our sequencing data. Our findings revealed here offer not only a rich genetic resource for further investigation of the functional roles of lncRNAs involved in *A. m. ligustica* response to *N. ceranae* infection, but also a novel insight into understanding host-pathogen interaction during microsporidiosis of honeybee.

## 1. Introduction

Honeybees are pivotal pollinators of crops and wild flora, and of great importance in supporting critical ecosystem balance [1]. In addition, honeybees serve as key models for studies on development, social behavior, disease transmission, and host-pathogen interaction [2]. Western honeybee (*Apis mellifera*) has been domesticated for honey production and crop pollination all over the world. The genome of *A. mellifera* was published in 2006 [3], which laid a solid foundation for its molecular and functional genomics studies. *Apis mellifera ligustica*, a subspecies of *A. mellifera*, is widely used in beekeeping industry in China and many other countries.

Microsporidia are spore-forming and obligate intracellular fungal pathogens, which can infect a wide variety of hosts such as mammals and insects [4]. Microsporidia infection occurs through the ingestion of spores in contaminated food or water, followed by the germination of these spores activated by the physical and chemical conditions inside the midgut; subsequently, the sporoplasm is injected into the host midgut epithelium cell where it multiplies; ultimately, the spores are excreted from the host in the feces, offering new sources of the infection through cleaning and feeding activities inside the colonies, or are disseminated into the environment [5–7]. Nosemosis is a serious disease in adult honeybees due to infection with *Nosema* species including *Nosema apis* and *Nosema ceranae*. The latter is a widespread microsporidian pathogen of honeybees, which was first identified by Fries et al. from *A. cerana* near Beijing [8], China, and shortly afterward, it was reported to spread to Europe [9] and Taiwan [10]. Currently, *N. ceranae* has been found in colonies of western honeybee throughout the world [11,12]. *N. ceranae* is infective to all castes in the colony [7], and it could dramatically reduce colony strength and productivity [13] as well as interacts with other environmental stressors to severely weaken colony health [14].

Previous studies revealed that much of the genome is transcribed, but only a small part of sequences have protein-coding capacity [15]. In human, only less than 2% of the genome contains conserved sequences for proteins [15]. Hence, many of the transcribed sequences in the genome are believed to be non-coding RNAs (ncRNAs), which are arbitrarily categorized into two types according to their sizes, one is small RNAs shorter than 200 nt, such as microRNAs (miRNAs) and small nucleolar RNAs (snoRNAs); and the other type is long non-coding RNAs (lncRNAs), which are longer than 200 nt and lack protein-coding potential [16]. LncRNAs can be further classified into antisense lncRNAs, intronic lncRNAs, overlapping lncRNAs, and intergenic lncRNAs [17]. LncRNAs are usually expressed at low levels, lack conservation among species, and often display tissue- or cell-specific expression patterns [18,19]. In recent years, lncRNAs have been found to play key roles in various biological processes in mammals and plants as potent regulators, natural miRNA target mimics, chromatin modifiers, and molecular cargo for protein re-localization [20]. Additionally, lncRNAs have been found to be closely related to some diseases such as Alzheimer’s disease [21] and acquired immune deficiency syndrome (AIDS) [22], therefore can be used as novel biomarkers and therapeutic targets. With the rapid development of high-throughput sequencing techniques, genome-wide investigations for lncRNAs have been conducted via cDNA/EST *in silico* mining [23,24], whole genome tilling array [25], and RNA-seq approaches [26]. By using deep sequencing and bioinformatics, more than 8,000 lncRNAs have been predicted in humans [18] and about 4,000 lncRNAs have been identified in mice [27,28]. In plants, 6,480 transcripts have been classified as lncRNAs in *Arabidopsis* [29]; 125 putative stress-responsive lncRNAs have been identified in wheat [30]. In microorganisms, our research group identified 379 novel lncRNAs in *Ascosphaera apis* (another common fungal pathogen of honeybee), 83 in *N. ceranae*, and revealed that these fungal lncRNAs shared similar characteristics with those in mammals and plants, such as shorter length and fewer exon number [31,32]. Recently, a number of lncRNAs were identified in insects such as *Plutella xylostella* [33], *Anopheles gambiae* [34], and *Bombycis mori* [35]. However, compared with mammals and plants, knowledge of honeybee lncRNA is still largely unknown. Thus far, only few lncRNAs have been discovered in honeybee, such as *lncov1*, *lncov2*, and *Ks-1* [36,37]. Utilizing transcriptome sequencing, Jayakodi et al. [38] identified 1514 long intergenic non-coding RNAs (lincRNAs) in *A. mellifera* and 2470 lincRNAs in *Apis cerana*, most of which had a tissue-specific expression pattern. More recently, Chen et al. [39] predicted a variety of lncRNAs, miRNAs and mRNAs during the ovary activation, oviposition inhibition and oviposition recovery processes; and they further found 73 differentially expressed genes and 14 differentially expressed lncRNAs (DElncRNAs) located in the QTL region, which may be candidate genes responsible for ovary size and oviposition.

To our knowledge, no study on honeybee lncRNAs response to fungal stress was reported until now, and understanding of potential roles of host stress-responsive lncRNAs was extremely limited. Here, to systematically identify lncRNAs, corresponding regulatory networks, and their potential roles involved in *A. m. ligustica* response to *N. ceranae* stress, we first performed whole transcriptome strand-specific RNA sequencing of normal and *N. ceranae*-stressed midgut samples of *A. m. ligustica* workers. We examined the expression pattern of host lncRNAs responding to *N. ceranae* challenge, followed by molecular validation of differentially expressed DElncRNAs. Moreover, regulatory networks of *A. m. ligustica* DElncRNAs were constructed and analyzed to further explore their potential roles during the fungal stress response. The current work generated a comprehensive list of *A. m. ligustica* lncRNAs, which will be a valuable complement to the other ncRNAs that have already been discovered in this important social insect. The results not only lay a foundation for deciphering the molecular mechanisms underlying *A. m. ligustica* response to *N. ceranae* stress, but also offer a beneficial resource for functional study on key *N. ceranae*-responsive lncRNAs in the future. Our data can also help better understanding the western honeybee-microsporidia interactions.

## 2. Materials and Methods

### 2.1. N. ceranae spore purification

Fresh spores were isolated from naturally-infected foragers from a colony located at Fuzhou city, Fujian province, China, following the method described by Cornman et al. [40] with some modifications [32]. (1) bees were kept in −20 °C for 5 min to anesthetize them, followed by separation of midguts with clean dissection tweezers, homogenization in distilled water, filtration in four layers of sterile gauze, and then three times of centrifugation at 8000 rpm for 5 min; (2) the supernatant was discarded as the spores remained in the sediment, the re-suspended pellet was further purified on a discontinuous Percoll gradient (Solarbio) consisting of 5 mL each of 25%, 50%, 75% and 100% Percoll solution, the spore suspension was overlaid onto the gradient and centrifuged at 14000 rpm for 90 min at 4 °C; (3) the spore pellet was carefully extracted with a sterile syringe and then centrifuged again on a discontinuous Percoll gradient to obtain clean spores (Figure S1A), which were frozen in liquid nitrogen and stored at −80 °C until deep sequencing, RT-PCR, and real-time quantitative PCR (RT-qPCR). A bit of spores were subjected to PCR identification and confirmed to be mono-specific using previously described primers [12]. The spore concentration was determined by counting using a CL kurt counter (JIMBIO) and the suspension was freshly prepared before use.

### 2.2. Experimental design and sample collection

Frames of sealed brood obtained from a healthy colony of *A. m. ligustica* located in the teaching apiary of College of Bee Science in Fujian Agriculture and Forestry University were kept in an incubator at 34 ± 0.5 °C, 50% RH to provide newly emerged *Nosema*-free honeybees. The emergent workers were carefully removed, confined to cages in groups of 20, and kept in the incubator at 32 ± 0.5 °C, 50% RH. The bees were fed *ad libitum* with a solution of sucrose (50% w/w in water). One day after eclosion, the honeybees were starved for 2 h and 20 workers per group were each immobilized and then fed with 5 μL of 50% sucrose solution containing 1×10^6^ spores of *N. ceranae* (Figure S1B). Those individuals that did not consume the total amount of solution were discarded from the assay. After feeding, bees were isolated for 30 min in individual vials in the growth chamber to ensure that the sugar solution was not transferred among honeybees and the entire dosage was ingested. Control bees were inoculated in an identical manner using a 50% sucrose solution (w/w in water) without *N. ceranae* spores. Three replicate cages of 20 honeybees each were used in *N. ceranae*-treated and untreated groups. Each cage was checked every 24 h and any dead bees removed. *N. ceranae*-treated and untreated workers’ midguts were respectively harvested 7 or 10 d post inoculation (dpi), immediately frozen in liquid nitrogen and kept at −80 °C until high-throughput sequencing and molecular experiments. treatment groups 7 and 10 dpi with sucrose solution containing *N. ceranae* spores were termed as Am7T (Am7T-1, Am7T-2, Am7T-3) and Am10T (Am10T-1, Am10T-2, Am10T-3); control groups 7 and 10 dpi with sucrose solution without *N. ceranae* spores were termed as Am7CK (Am7CK-1, Am7CK-2, Am7CK-3) and Am10CK (Am10CK-1, Am10CK-2, Am10CK-3).

### 2.3. RNA extraction, strand-specific cDNA library construction and deep sequencing

Total RNA of the six midgut samples from *N. ceranae*-treated groups and six midgut samples from untreated groups were respectively extracted usng Trizol (Life Technologies) following the manufacturer’s instructions, and checked via 1% agarose gel eletrophoresis. Subsequently, rRNAs were removed to retain mRNAs and ncRNAs, which were fragmented into short fragments by using fragmentation buffer (Illumina) and reverse transcripted into cDNA with random primers. Next, second-strand cDNA were synthesized by DNA polymerase I, RNase H, dNTP (dUTP instead of dTTP), and buffer. The cDNA fragments were purified using QiaQuick PCR extraction kit (QIAGEN), end repaired, poly(A) added, and ligated to Illumina sequencing adapters. UNG (Uracil-N-Glycosylase) (Illumina) was then used to digest the second-strand cDNA. Ultimately, the digested products were size selected by agarose gel electrophoresis, PCR amplified, and sequenced on Illumina HiSeq™ 4000 platform (Illumina) by Gene Denovo Biotechnology Co. (Guangzhou). All RNA sequencing data produced in our study are available in NCBI Short Read Archive database (http://www.ncbi.nlm.nih.gov/sra/) and could be accessed under the SRA accession number: SUB3038297.

### 2.4. Quality control and mapping of reads

Reads produced from the sequencing machines included raw reads containing adapters or low quality bases which would affect the following assembly and analysis. Therefore, reads were further filtered by removing reads containing adapters, more than 10% of unknown nucleotides (N), and more than 50% of low quality (*Q*-value ≤ 20) bases to gain high quality clean reads.

Short reads alignment tool Bowtie2 [41] was used for mapping reads to ribosome RNA (rRNA) database. The mapped reads were then removed and the remaining reads were further used in assembly and analysis of transcriptome. The rRNA removed reads of each sample were then mapped to reference genome of *Apis mellifera* (assembly Amel_4.5) by TopHat2 (version 2.0.3.12) [42]. The alignment parameters were as follows: (1) maximum read mismatch is two; (2) the distance between mate-pair reads is 50 bp; (3) the error of distance between mate-pair reads is ± 80 bp.

### 2.5. Transcripts assembly

Transcripts were assembled using software Cufflinks [43], which together with TopHat2, allow researchers to identify novel genes and novel splice variants of known ones. The program reference annotation based transcripts (RABT) was preferred. Cufflinks constructed faux reads according to reference to make up for the influence of low coverage sequencing. During the last step of assembly, all of the reassembles fragments were aligned with reference genes and then similar fragments were removed. Cuffmerge was used to merge transcripts from different replicas of a group into a comprehensive set of transcripts, and the transcripts from multiple groups were then merged into a finally comprehensive set of transcripts for further downstream analyses.

### 2.6. Bioinformatic pipeline for identification and annotation of lncRNAs, and quantification

To identify the novel transcripts, all of the reconstructed transcripts were aligned to reference genome of *Apis mellifera* (assembly Amel_4.5) and were divided into twelve categories by using Cuffcompare [43]. Transcripts with one of the classcodes “u, i, j, x, c, e, o” were defined as novel transcripts. The following parameters were used to identify reliable novel lncRNAs: the length of transcript was longer than 200 bp and the exon number was more than two; novel transcripts were then aligned to the Nr, GO (Gene Ontology) and KEGG (Kyoto Encyclopedia of Genes and Genomes) databases to obtain protein functional annotation. Softwares CNCI [44] and CPC [45] were utilized in combination to sort non-protein coding RNA candidates from putative protein-coding RNAs in the unknown transcripts by default parameters. The intersection of both results was chosen as lncRNAs. The different types of lncRNAs including lincRNA, intronic lncRNA, anti-sense lncRNA were selected using cuffcompare. The detailed flow of novel lncRNA prediction is shown in Figure S2.

Transcripts abundances were quantified by software RSEM [46] following (1) a set of reference transcript sequences were generated, preprocessed according to known transcripts, new transcripts (in FASTA format), and gene annotation files (in GTF format); (2) reads were realigned to the reference transcripts by Bowtie alignment program and the resulting alignments were used to estimate transcript abundances.

The transcript expression level was normalized by using FPKM (fragments per kilobase of transcript per million mapped reads) method, which can eliminate the influence of different transcripts lengths and sequencing data amount on the calculation of transcripts expression. Therefore, the calculated transcripts expression can be directly used for comparing the difference of transcripts expression among samples.

### 2.7. DElncRNAs, target gene, and ceRNA analyses

DElncRNAs between any two libraries were identified by edgeR [47] (release 3.2). The thresholds used to evaluate the statistical significance of differences in lncRNA expression were defined as FDR < 0.05 and an absolute value of the log2 (Fold change) > 1.

*Cis*-acting lncRNAs function via targeting neighbouring genes [48,49]. In the present study, we searched for coding genes in the regions located 10-kb upstream and downstream of all of the identified lncRNAs for predicting their functional roles.

All neighbouring genes were mapped to GO terms in the GO database (http://www.geneontology.org/), and gene numbers were calculated for each term; significantly enriched GO terms in neighbouring genes comparing to the reference genome background were defined by hypergeometric test. KEGG pathway enrichment analysis was conducted using KOBAS 2.0, with the *A. mellifera* genome as background. Only GO terms or KEGG pathways with corrected *p*-values of less than 0.05 were considered enriched.

Following traditional miRNA target prediction methods, we inferred the conserved regions of *A. m. ligustica* lncRNAs that may harbor MREs for ceRNA networks. miRanda(v3.3a) [50] (animal), RNAhybrid(v2.1.2)+svm_light(v6.01) [51,52] (animal) and TargetFinder(Version: 7.0) [53] (plant) were used to predict MREs in the conserved regions of lncRNAs.

### 2.8. Real-time quantitative PCR (RT-qPCR) confirmation of DElncRNAs

The first cDNA strand was synthesized using the SuperScript first-strand synthesis system (TaKaRa) according to the protocol. Primers for qPCR were designed using the DNAMAN software and synthesized by Sangon Biotech Co., Ltd. The housekeeping gene *actin* was used as an internal control. The RNA samples used as templates for RNA-seq were the same as those used for RT-qPCR, which was conducted on a QuanStudio Real-Time PCR System (ThemoFisher). The 20 μL PCR reaction mixture contained 10 μL SYBR Green dye (Vazyme); 0.4 μL (10 pmol/μL) specific forward primer; 0.4 μL (10 pmol/μL) reverse primer; 0.4 μL ROX reference dye; 2 μL (10 ng/μL) diluted cDNA; and 6.8 μL RNase free water. Cycling parameters were as follows: 95 °C for 1 min, followed by 40 cycles of 95 °C for 15 s, 60 °C for 30 s, and 72 °C for 45 s. The relative changes of selected gene expression was calculated using the 2^−ΔΔCT^ method [54]. These assays were performed in triplicate. The specific primers used in RT-qPCR are shown in Table S1.

### 2.9. Statistical analysis

All statistical analyses were performed using SPSS software (IBM) and GraphPad Prism 6.0 software (GraphPad Software Inc.). Data were presented as mean ± standard deviation (SD). Statistical analysis was calculated using independent-samples t-test and one way ANOVA. Fisher’s exact test was employed to filter the significant GO terms and KEGG pathways using R software 3.3.1. *P* < 0.05 was considered statistically significant.

## 3. Results

### 3.1. Sequencing results and quality control

In our study, a total of 1956129858 raw reads were produced from 12 cDNA libraries, and 1946489304 clean reads were obtained after strict quality control (Table 1). The percentage of clean reads among raw reads in each library ranged from 99.42 to 99.57%, with a mean Q30 of 93.82% (Table 1). In addition, clean reads were aligned with the reference genome of *A. mellifera*, and the result showed mapping ratio of 12 samples ranged from 39.48 to 60.76%. Among these mapped reads, 66.17–70.07% were mapped to coding DNA sequence regions, 5.34–8.18% to intron regions, 14.49–16.08% to intergenic regions, and 8.83–11.00% to untranslated regions. Moreover, high Pearson correlation coefficients (0.9119-0.9993) were found among biological replicates of each group, suggesting the reproducibility of sample preparation (Figure S3).

**Table 1.**
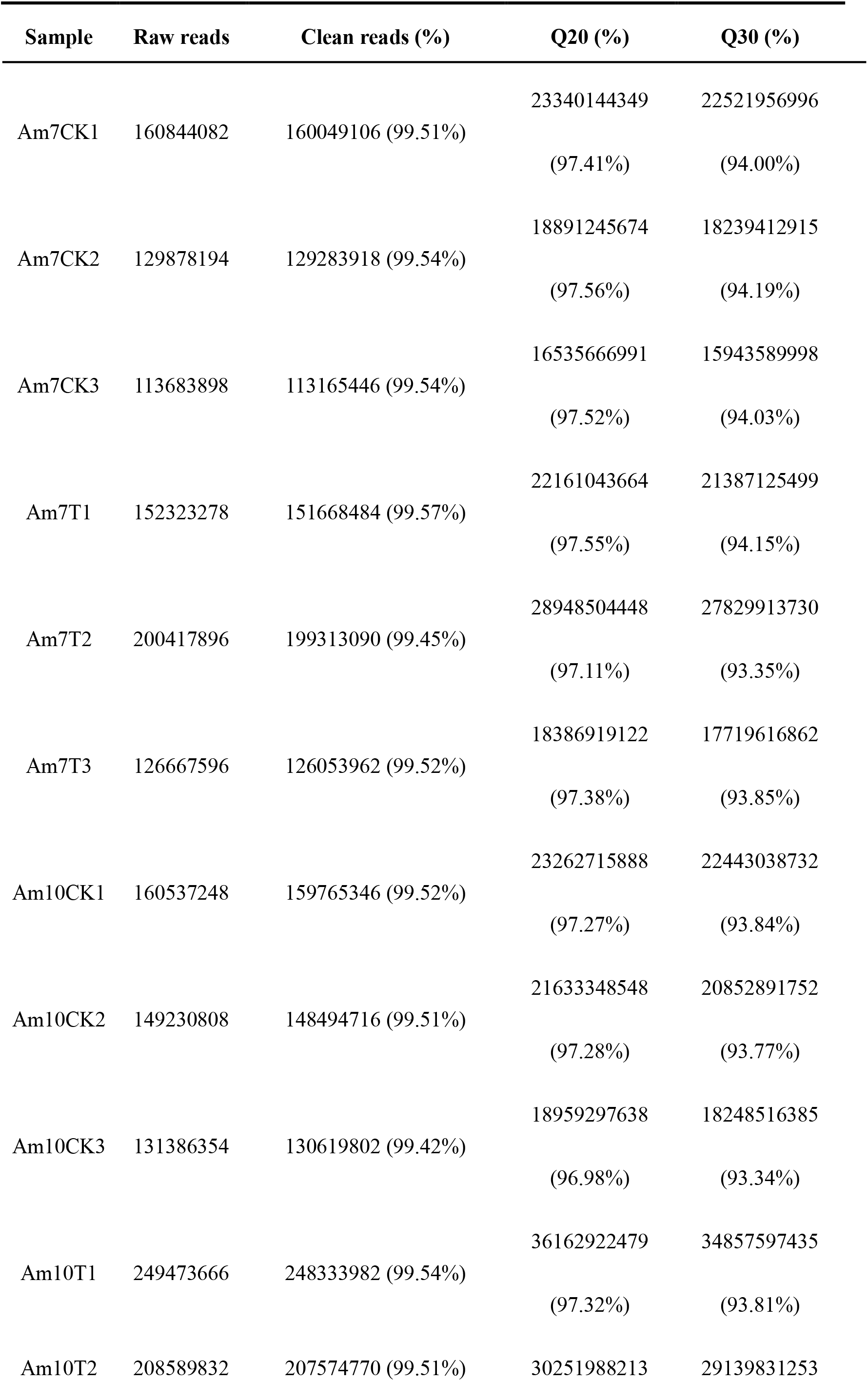
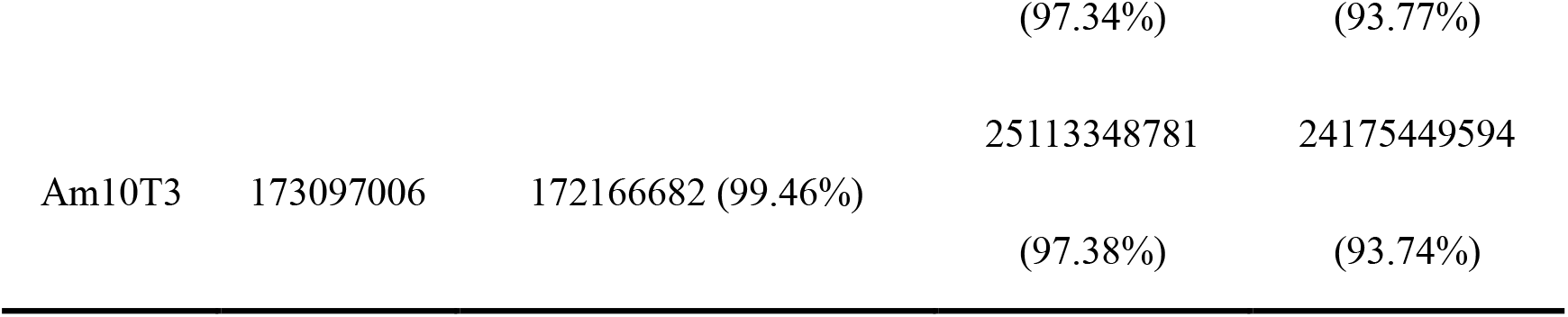
Quality control of transcriptome data.

### 3.2. Characterization and validation of A. m. ligustica lncRNAs

A high stringency filtering process (Figure S2) was used to remove low quality lncRNA transcripts. In total, 6353 lncRNAs were identified from midgut samples, including 4749 known lncRNAs and 1604 novel lncRNAs. These *A. m. ligustica* lncRNAs were found to be shorter in exon and intron length and fewer in exon number than protein-coding genes (Figure 1A-C), which is in accordance with findings in previous studies [55–59]. Additionally, the expression level of each transcript was estimated and the result showed the levels of lncRNAs are lower than those of mRNAs in the midgut of *A. m. ligustica* worker (Figure 1D).

**Figure 1.**
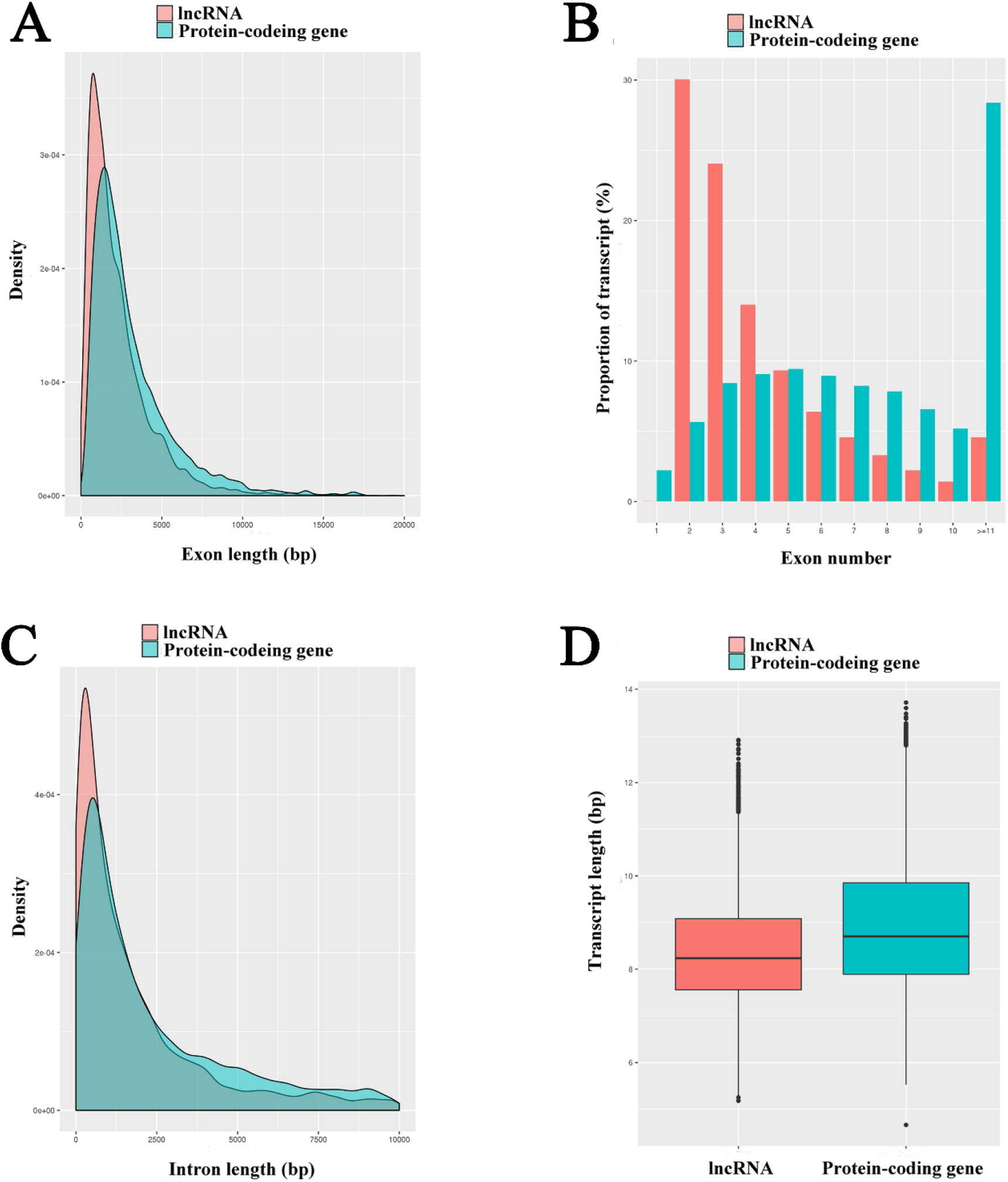
Properties of *A. m. ligustica* lncRNAs. **(A)** Exon size distributions for lncRNAs and protein-coding transcripts. **(B)** Number of exons per lncRNAs and protein-coding transcripts. **(C)** Intron size distributions for lncRNAs and protein-coding transcripts. **(D)** Expression levels of lncRNAs and protein-coding transcripts.

### 3.3. Identification of A. m. ligustica lncRNAs that respond to N. ceranae stress

Our main objective was to identify candidate *N. ceranae*-responsive lncRNAs involved in *A. m. ligustica* workers’ midguts. In total, 111 lncRNAs were differentially expressed in Am7CK vs Am7T, including 62 up-regulated and 49 down-regulated ones (Figure 2A, Table S2); while in Am10CK vs Am10T, 146 DElncRNAs including 82 up-regulated and 64 down-regulated ones were identified (Figure 2A, Table S3). The expression clustering of DElncRNAs in Am7CK vs Am7T and Am10CK vs Am10T was further conducted, and the result showed various DElncRNAs have differential expression levels (Figure 2B-E). Among them, TCONS_00037745 and TCONS_00029069 was the most up-regulated, while XR_001706167.1 and TCONS_00011956 was the most down-regulated. In addition, 857 and 971 mRNAs showing differential expressions in *N. ceranae*-treated groups compared with control groups (472 up-regulated and 385 down-regulated mRNAs in Am7CK vs Am7T, 611 up-regulated and 360 down-regulated mRNAs in Am10CK vs Am10T) (Figure 2A). Similar differential expression trends of mRNAs were observed in the volcano plots (Figure 2F-G). The number of DEGs and DElncRNAs among two comparison groups demonstrated an increase as the *N. ceranae* stress progressed (Figure 2A).

**Figure 2.**
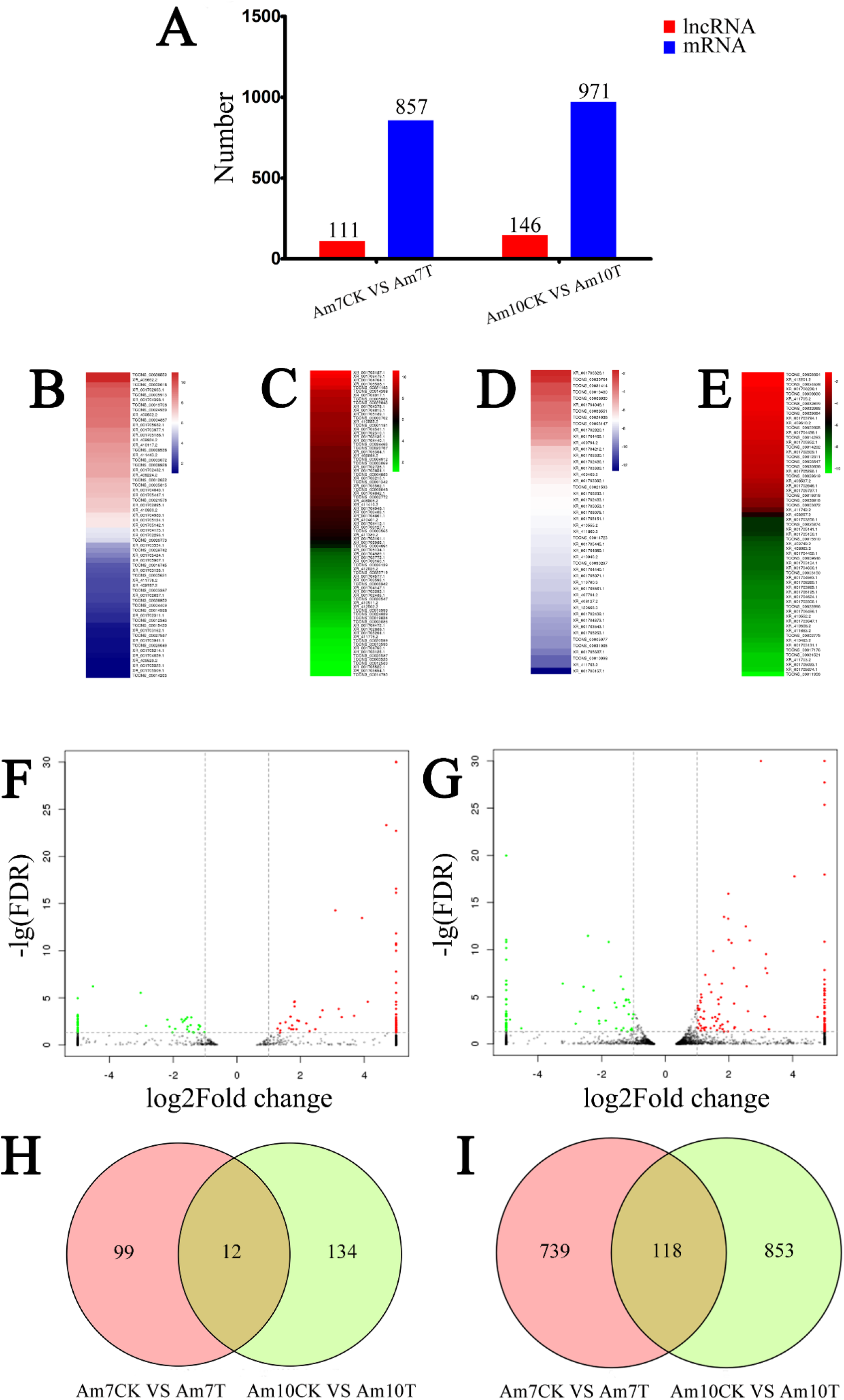
Differential expression patterns of *A. m. ligustica* lncRNAs and mRNAs in *N. ceranae*-stressed midguts compared with normal midguts. **(A)** Number of DElncRNAs and DEGs. **(B-C)** Expression clustering of up- and down-regulated lncRNAs in Am7CK vs Am7T. **(D-E)** Expression clustering of up- and down-regulated lncRNAs in Am10CK vs Am10T. **(F-G)** Volcano plots of DEGs in Am7CK vs Am7T and Am10CK vs Am10T. **(H)** Venn diagram of DElncRNAs in Am7CK vs Am7T and Am10CK vs Am10T. **(I)** Venn diagram of DEGs in Am7CK vs Am7T and Am10CK vs Am10T.

Moreover, Venn analyses showed 12 DElncRNAs were shared by Am7CK vs Am7T and Am10CK vs Am10T, while the numbers of unique DElncRNAs in the two comparison groups were 99 and 134, respectively (Figure 2H); 118 DEGs were common between Am7CK vs Am7T and Am10CK vs Am10T, while 739 and 853 were specific in the two comparison groups, respectively (Figure 2I)

### 3.4. Functional investigation of N. ceranae-responsive lncRNAs in A. m. ligustica workers’ midguts

Previous studies proved that lncRNAs could regulate the expression of target genes via chromatin remodeling, control of transcription initiation, and post-transcriptional processing [60,61]. LncRNAs can regulate target gene expression by acting in *cis* on neighbouring loci [60]. In the current work, we investigated the *cis* role of lncRNAs by screening the protein-coding genes as potential lncRNA *cis*-regulatory targets in the regions located 10-kb upstream and downstream of all the identified lncRNAs for prediction of their functional roles. GO analyses suggested the putative target genes of DElncRNAs in Am7CK vs Am7T were annotated as 10 biological process-associated terms such as metabolic process (17) and cellular process (17), 10 molecular function-related terms such as binding (26) and catalytic activity (23), 11 cellular component-connected terms such as cell (8) and membrane (6); similarly, the targets of DElncRNAs in Am10CK vs Am10T were annotated with 14 biological process-associated terms (eg. single-organism process, localization, and biological regulation), 11 molecular function-related terms (eg. transporter activity, signal transducer activity, and molecular transducer activity), and 13 cellular component-connected terms (eg. organelle, macromolecular complex, and extracellular region). The top 15 significant GO terms were respectively presented in Table S4 and Table S5.

KEGG pathway enrichment analyses concluded that 27 neighboring genes of DElncRNAs in Am7CK vs Am7T were enriched in 47 pathways associated with organismal systems (seven; eg. immune system and aging), metabolism (22; eg. carbon metabolism and purine metabolism), genetic information processing (10; eg. DNA replication and ribosome), environmental information processing (four; eg. signal transduction and membrane transport), and cellular processes (peroxisome and lysosome); comparatively, 41 neighboring genes of DElncRNAs in Am10CK vs Am10T were enriched in 50 pathways, including 30 material and energy related ones such as starch and sucrose metabolism (TCONS_00010783, XM_394494.6), and glycerolipid metabolism (XM_006562913.2), and cellular pathways such as ubiquitin mediated proteolysis (TCONS_00027323, XM_006570714.2), endocytosis (XM_016915026.1, XM_016917619.1), and lysosome (XM_016914010.1). The top 15 significant enriched pathways were respectively shown in Table S6 and Table S7.

### 3.5. Discovery of A. m. ligustica lncRNAs as miRNA precursors and ceRNAs

LncRNA loci that overlapped with miRNA loci on the same strand were regarded as the miRNA precursors [62]. To determine whether lncRNAs are in fact precursors of miRNAs, the lncRNA sequences were compared with the miRNA sequences that obtained from miRBase. The result showed that 27 lncRNAs harbored eight complete known miRNA precursors (Table S8); in addition, the secondary structures of lncRNA transcripts suggested many known and novel lncRNAs contained a stable hairpin structure for miRNA precursors. For example, TCONS_00019779 harbored ame-mir-927a (Figure S4); TCONS_00036128 harbored ame-mir-1-1 and ame-mir-750. Additionally, another 513 *A. m. ligustica* lncRNAs were predicted to be precursors of 2257 novel miRNAs (Table S9).

The competitive endogenous RNAs (ceRNAs) including mRNAs and long noncoding RNAs (lncRNAs) contain shared miRNA response elements (MREs), and they can compete for miRNA binding [63]. LncRNAs may bind miRNAs as ceRNAs, thereby functioning as miRNA sponges [64]. The lncRNA-miRNA interaction can be examined using traditional miRNA target prediction methods [65,66]. Here, we analyzed the 6,353 lncRNA transcripts that may harbor MREs for ceRNA network [63] using miRanda [50] PITA [67], and RNAhybrid [51,52]. As shown in Figure 3, complex ceRNA networks of *A. m. ligustica* DElncRNAs and their target miRNAs were visualized using Cytoscape. A total of 106 DElncRNAs in Am7CK vs Am7T were detected to target 76 *A. m. ligustica* miRNAs (Table S10). Of these, some DElncRNAs were targeted by more than one miRNAs. For example, XR_001702296.1 and TCONS_00030779 could be targeted by 22 and 19 miRNAs, respectively; additionally, some DElncRNAs were targeted by only one miRNA, such as XR_409934.2, XR_409794.2, and XR_001703543.1. Meanwhile, several lncRNAs had the same target miRNA. For example, XR_410555.2, XR_001705522.1, and TCONS_00030779 can target mir-941-y; as many as 23 lncRNAs including XR_001703554.1 and TCONS_00031414 could target novel-m0007-5p. In Am10CK vs Am10T comparison group, 143 DElncRNAs were predicted to be targets of 98 miRNAs (Table S11). Similarly, part of DElncRNAs such as XR_412201.2 and TCONS_00015510 were targeted by several miRNAs, while some (eg. XR_412502.2 and XR_409610.2) had only one target miRNA. In addition, some DElncRNAs including XR_001706086.1, XR_001702485.1, and TCONS_00036139 were targeted by the same miRNA (mir-9189-y).

**Figure 3.**
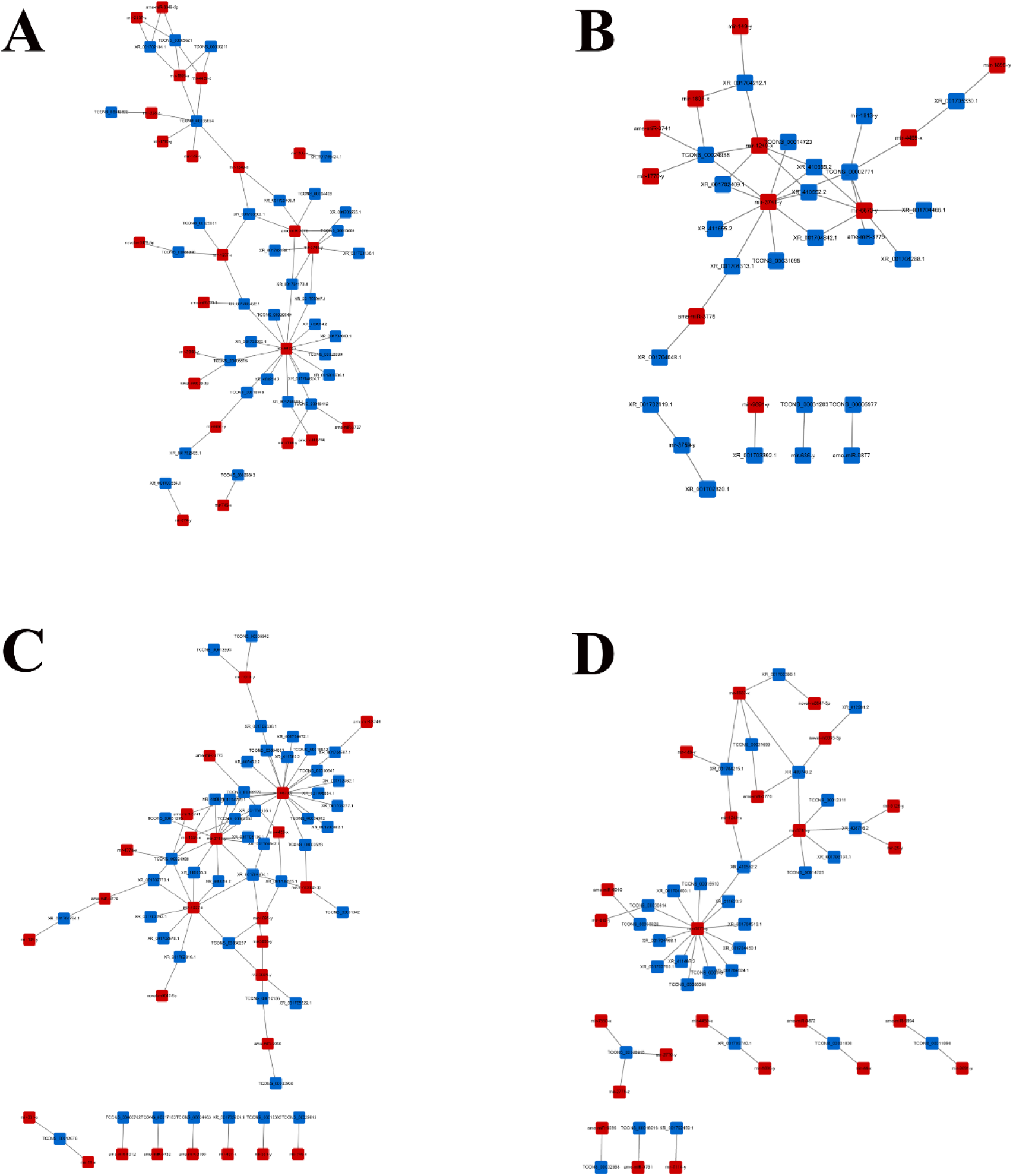
ceRNA networks of DElncRNAs in *N. ceranae*-stressed and normal midguts of *A. m. ligustica* workers. **(A-B)** ceRNA networks of up- and down-regulated lncRNAs in Am7CK vs Am7T. **(C-D)** ceRNA networks of up- and down-regulated lncRNAs in Am10CK vs Am10T.

### 3.6. DElncRNA-miRNA-mRNA regulatory networks in A. m. ligustica workers’ midguts invaded by N. ceranae

To further investigate the roles of DElncRNAs, target mRNAs of DElncRNA-targeted miRNAs were predicted using miRanda [50], RNAhybrid [51,52] and TargetFinder [53]. 278 and 365 target mRNAs were observed in Am7CK vs Am7T and Am10CK vs Am10T. DElncRNA-miRNA-mRNA regulatory networks were constructed with Cytoscape, and it’s discovered that DElncRNAs, target miRNAs of DElncRNAs, and target mRNAs of DElncRNA-targeted miRNAs formed even more complex networks (Figure 4). GO categorizations demonstrated that target genes in Am7CK vs Am7T were involved in 14 biological process-related terms including cellular process, metabolic process, and biological regulation; nine molecular function-related terms including binding, catalytic activity, and molecular function regulator; 10 cellular component-related terms including cell, membrane, and organelle (Figure 5A); while target genes in Am10CK vs Am10T were associated with 28 GO terms, which also include the above-mentioned ones (Figure 5B). Moreover, we found in Am7CK vs Am7T and Am10CK vs Am10T, 12 and 18 target genes were engaged in response to stimulus; 12 and 16 target genes were involved in signaling, respectively (Figure 5).

**Figure 4.**
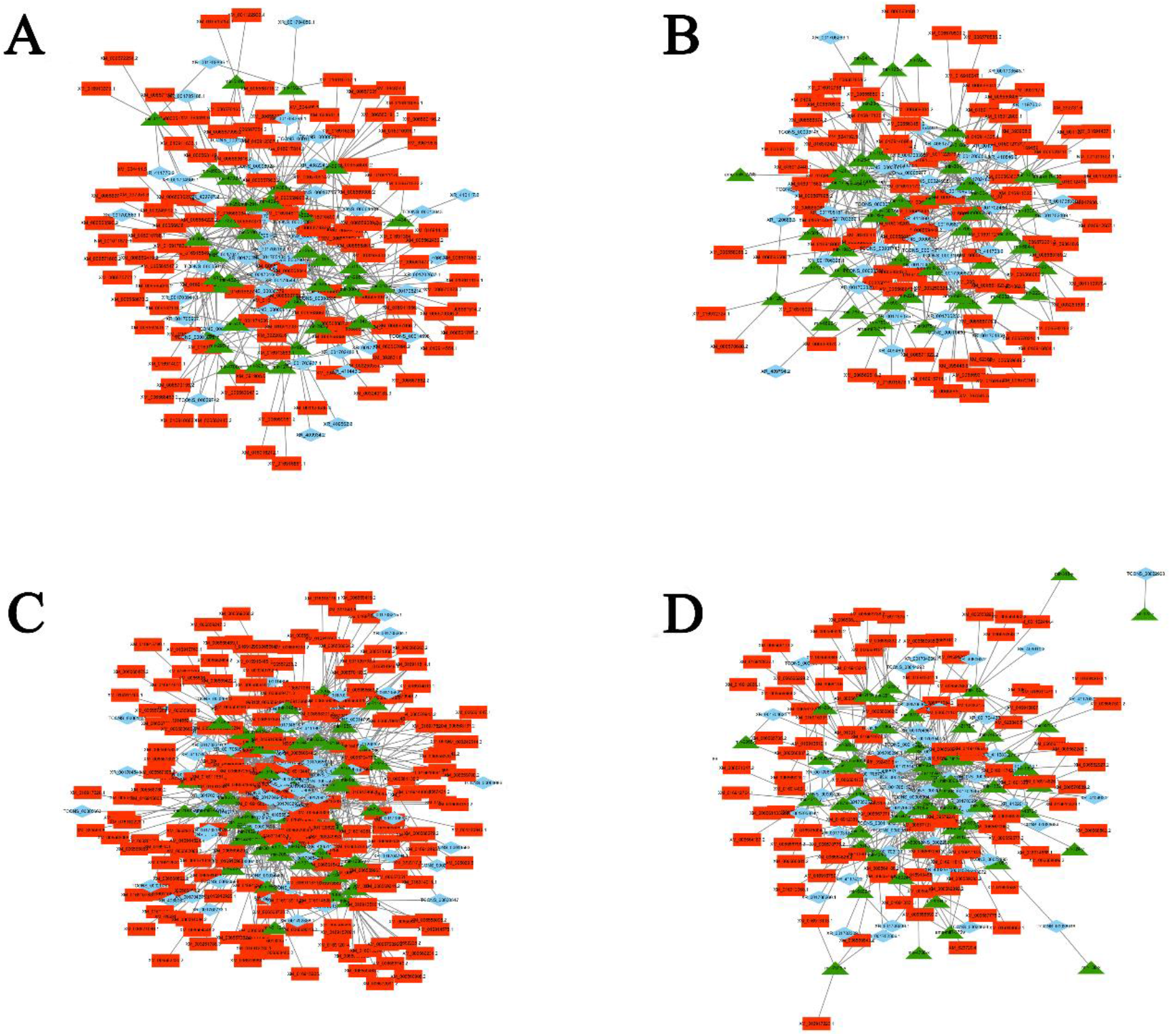
DElncRNA-miRNA-mRNA regulatory networks in *N. ceranae*-stressed and normal midguts of *A. m. ligustica* workers. **(A-B)** DElncRNA-miRNA-mRNA regulatory networks of up- and down-regulated lncRNAs in Am7CK vs Am7T. **(C-D)** DElncRNA-miRNA-mRNA regulatory networks of up- and down-regulated lncRNAs in Am10CK vs Am10T.

**Figure 5.**
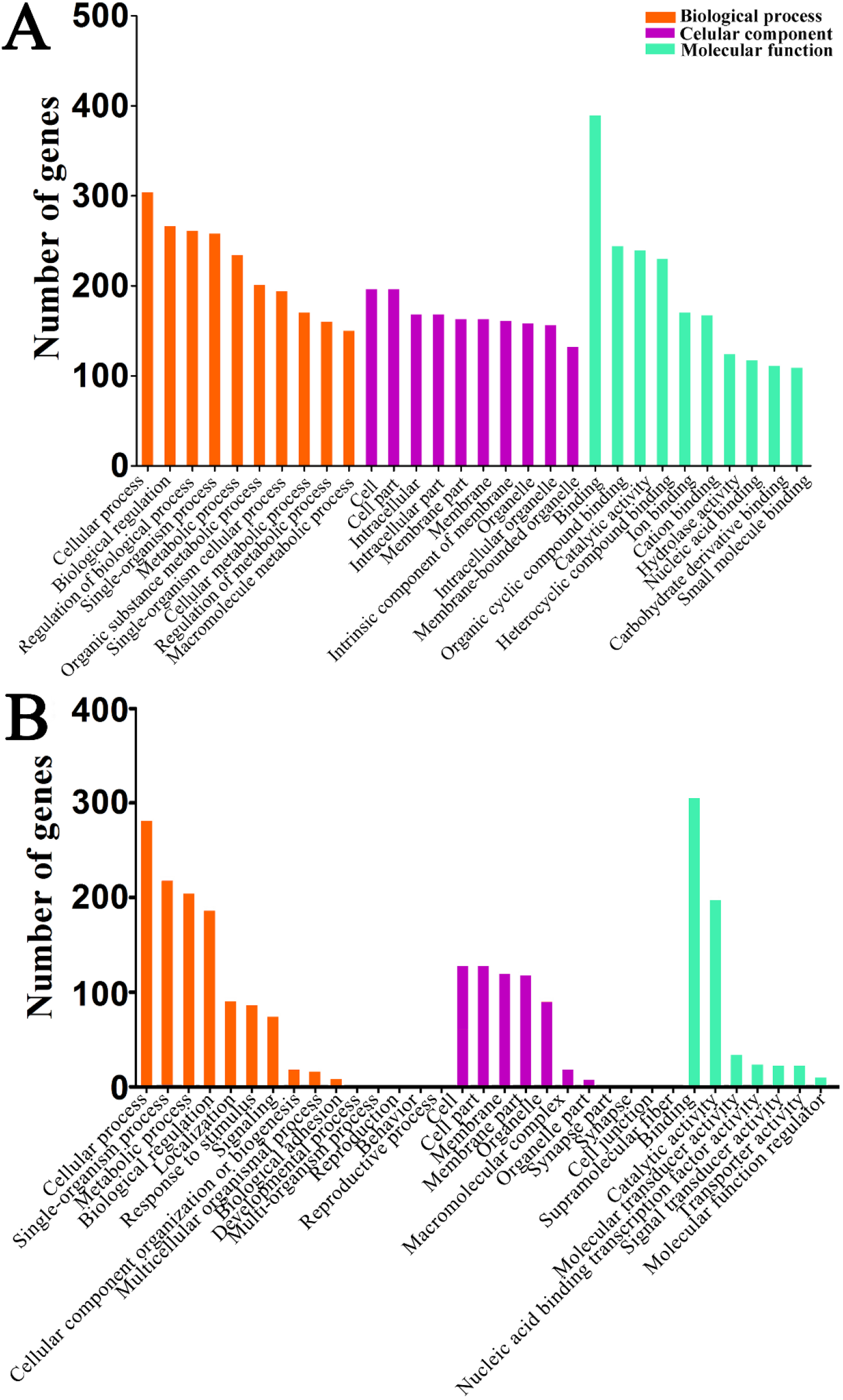
GO categorizations of target genes of DElncRNA-targeted miRNAs in Am7CK vs Am7T **(A)** and Am10CK vs Am10T **(B).**

Further, pathway analyses demonstrated target genes in Am7CK vs Am7T were enriched in 39 pathways, including 19 metabolism-related pathways such as biosynthesis of amino acids and oxidative phosphorylation, eight genetic information processing-related pathways such as transcription and translation; six signal transduction-related pathways such as Wnt signaling pathway and Hippo signaling pathway (Figure 6A); while target genes in Am10CK vs Am10T were involved in 45 pathways, among them 23, seven, and seven were relevant with metabolism, genetic information processing, and signal transduction, respectively (Figure 6B). Interestingly, target genes in both Am7CK vs Am7T and Am10CK vs Am10T were enriched in cellular immunity-related pathways including endocytosis, phagosome, and ubiquitin mediated proteolysis; however, only five target genes in Am7CK vs Am7T were associated with lysosome, no gene in Am10CK vs Am10T was found to be associated with any humoral immune pathway (Figure 6).

**Figure 6.**
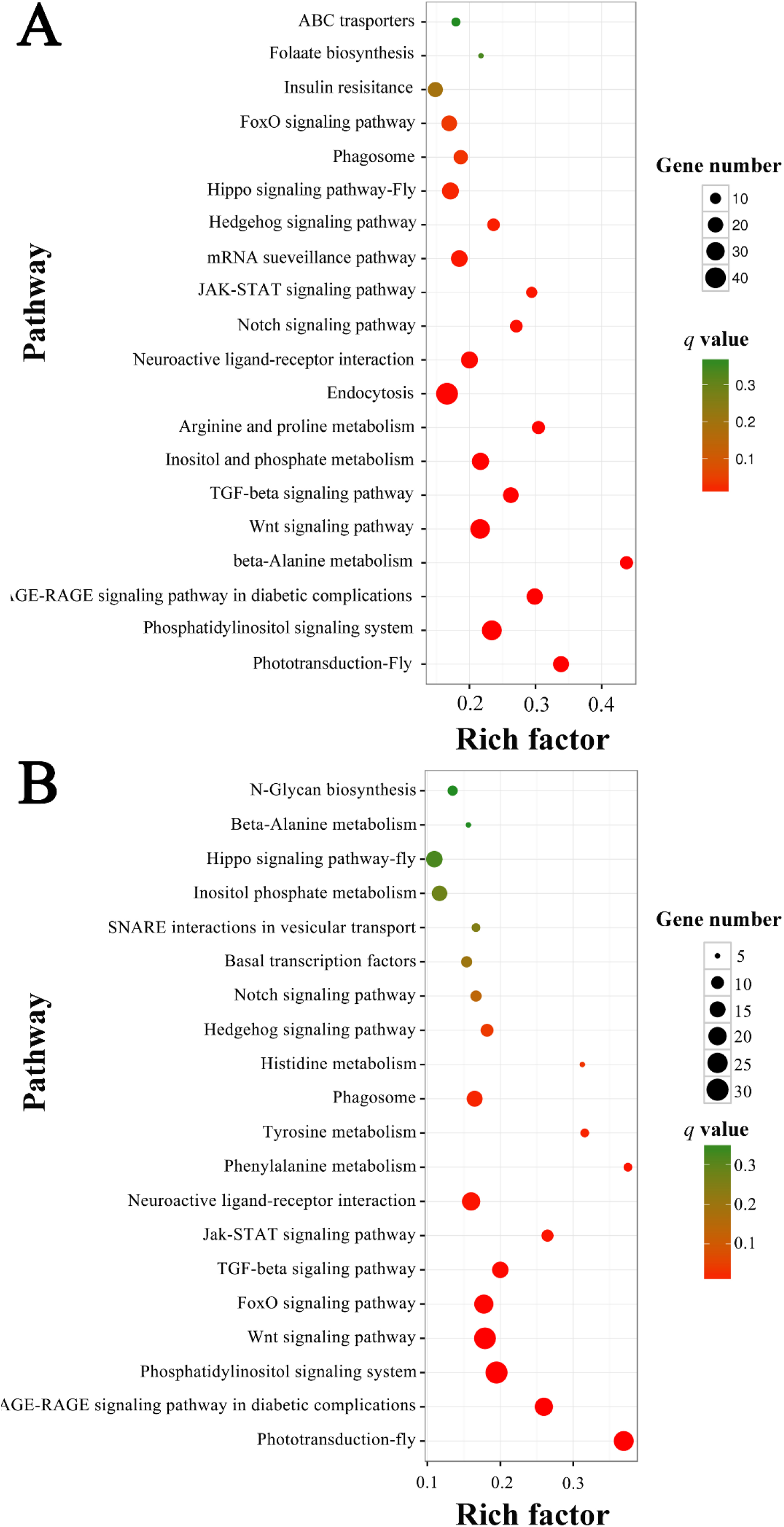
KEGG pathway enrichment analyses of target genes of DElncRNA-targeted miRNAs in Am7CK vs Am7T **(A)** and Am10CK vs Am10T **(B).**

### 3.6. Validation of DElncRNAs byRT-qPCR

To validate our RNA-seq data, 12 DElncRNAs were randomly selected for RT-qPCR assay, including six from Am7CK vs Am7T and six from Am10CK vs Am10T. The results showed that the expression patterns of 10 DElncRNAs were in agreement with the RNA-seq results (Figure 7), confirming our transcriptome sequencing data and analyses.

**Figure 7.**
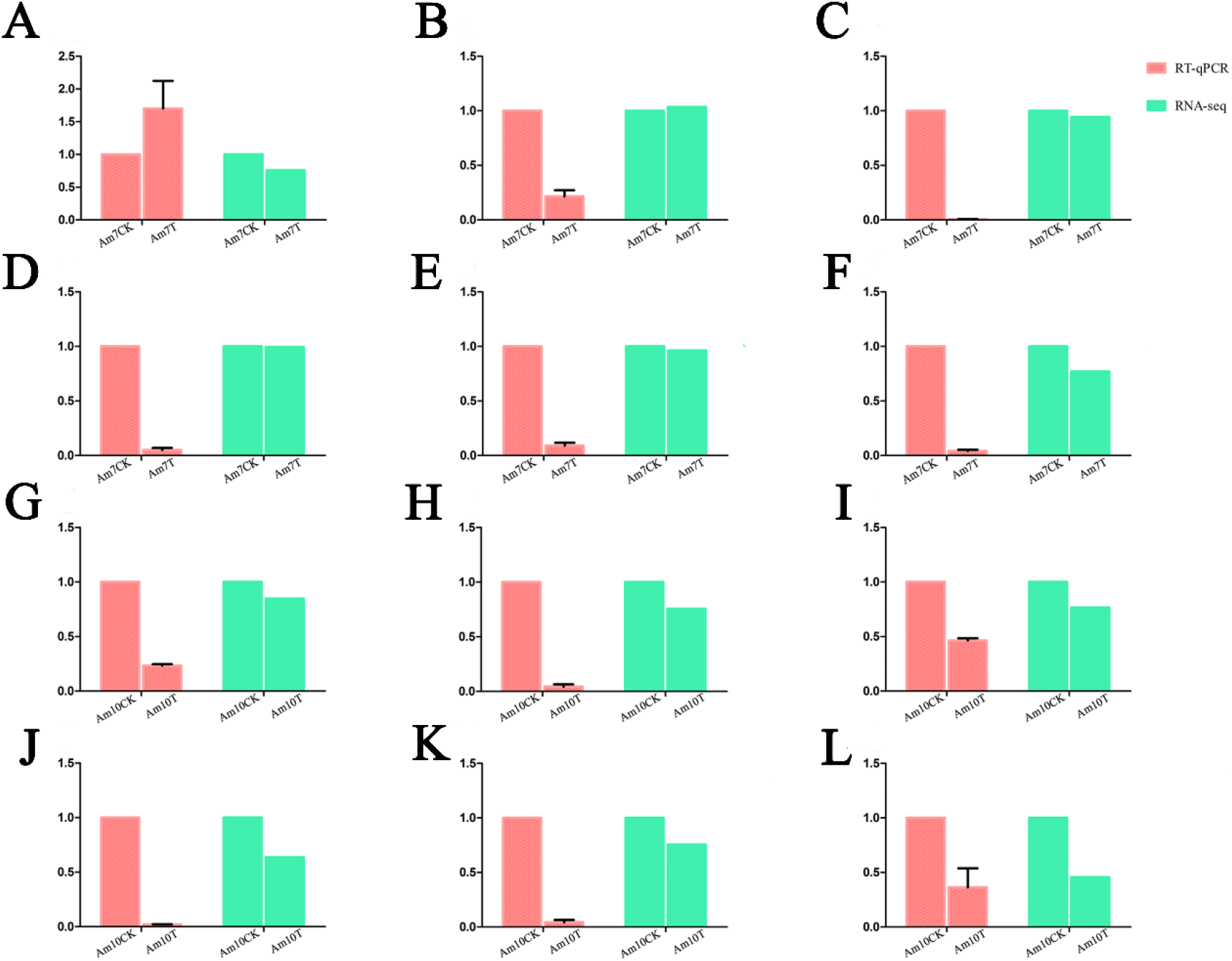
Validation of the expression patterns of *A. m. ligustica* lncRNAs via RT-qPCR **(A-L).**

## 4. Discussion

In the last decade, lncRNA has become a worldwide research hotspot attracting increasing attention, and the overwhelming majority of lncRNA studies have been performed in mammals and plants, especially in model species such as human [18] and *Arabidopsis* [29]. However, studies on insect lncRNAs are still in the initial stage. Recently, a series of lncRNAs has been discovered in several insect species, including *Drosophila melanogaster* [68], *Anopheles gambiae* [34], *Nilaparvata lugens* [69], and *Bombyx mori* [35]. Here, for the first time, we used rRNA removal and strand-specific RNA sequencing to systematically characterize and identify lncRNAs involved in responses of *A. m. ligustica* to *N. ceranae* infection. In comparison with polyA enrichment sequencing, this method has an obvious advantage of allowing non-polyA transcripts to be gained [70]; hence, strand information of lncRNAs was also included in our sequencing data, allowing to distinguishing sense transcripts from antisense transcripts. Considering different kinds of lncRNAs may play their parts in various manners, a detailed categorization of lncRNAs would facilitate further understanding their multiple functions [20]. In this work, 4749 conserved lncRNAs and 1604 novel lncRNAs were predicted from normal and *N. ceranae*-stressed midguts of *A. m. ligustica* workers, which offered a relatively robust list of potential lncRNAs for *A. m. ligustica*. This set of lncRNAs will be beneficial for functional genomics research and complementing the reference genome annotation of *Apis mellifera*. In our study, we detected that most lncRNAs were expressed at relatively low levels following an FPKM cutoff, thus lncRNAs with low expression may be ignored. Higher RNA-seq coverage can partly help overcome this problem [71]. Considering that lncRNAs are often expressed in a tissue- or development-specific manner [72], it’s believe that the identified lncRNAs just occupied a fraction of the total lncRNAs in *A. m. ligustica*, and more lncRNAs may be discovered using different castes, various organs and tissues, and organs and tissues under different stresses.

In our study, these *A. m. ligustica* lncRNAs were found to share some features with their counterparts in other species, including relatively short lengths, low exon numbers, and relatively low expression [73–75], indicating that these characteristics are common for lncRNAs in most species [34,35,68,69]. In vertebrates, lncRNAs have poor sequence conservation compared to protein-coding genes, making it quite difficult to predict the functions of lncRNAs simply based on their nucleotide sequences [17,18,76,77]. For example, less than 6% of zebrafish lncRNAs exhibited sequence conservation with human or mouse lncRNAs. Previous studies have demonstrated that the sequence conservation of lncRNAs between human and other species were only about 12% [78 79]. In the present study, the identified lncRNAs were compared against the NCBI Nr and NONCODE databases, but no highly similar sequences were found (data not shown), indicative of a lack of sequence conservation of *A. m. ligustica* lncRNAs.

Under abiotic and biotic stresses, lncRNAs could be used to regulate gene expression in multiple ways. For example, lncRNA plays pivotal roles in controlling the stress-response in plants including *Populus* [80] and wheat [30]. Jayakodi et al. identified 15 long intergenic non-coding RNAs (lincRNAs) showing significant differential expression between sacbrood virus (SBV)-infected and uninfected *Apis cerana*, and further confirmed the expression of 11 lincRNAs using RT-qPCR [38]. In this work, the expression levels of 111 and 146 lncRNAs were observed to significantly alter in *N. ceranae*-infected midguts of *A. m. ligustica* workers at 7 dpi and 10 dpi, which suggested the expression of some lncRNAs were affected by *N. ceranae* stress.

The functions of lncRNAs are highly diverse; however, only few known lncRNAs had functional annotations. To predict the functional roles of lncRNAs, their correlated protein coding genes and associated biological pathways are always surveyed to gain useful information [27]. The *cis* effect is defined as the regulatory action of lncRNAs on genes located upstream or downstream, which had been proved to be a common mechanism [81]. Here, to further investigate the roles of lncRNAs involved in *A. m. ligustica* response to *N. ceranae*, the potential function of the DElncRNAs was predicted using *cis* method. GO classifications suggested the targets of DElncRNAs in Am7CK vs Am7T and Am10CK vs Am10T were respectively involved in 31 and 38 functional terms, among them binding-associated activity was the abundant one in both comparison groups. LncRNAs are key regulators in various biological functions, but the related mechanisms are not fully understood. One of the regulatory mechanisms is based on interactions between different biological macromolecules such as RNA-RNA and RNA-DNA interactions [82]. Therefore, it’s believed that *A. m. ligustica* lncRNAs were likely to participate in *N. ceranae*-response via these interactions. However, it should be noticed that the evidence presented here are indirect, and further experiments are needed to verify the interaction among these lncRNAs and their targets. Further, pathway analyses demonstrated that the DElncRNAs were involved in regulating 47 and 50 pathways. *Nosema ceranae* is a mitochondriate that has a high dependency on host ATP, leading honeybee to increase sucrose needs and glycometabolism-relative genes expression [83,84]. Similarly, in this study, neighboring genes of DElncRNAs were enriched in multiple pathways associated with glucose metabolism, including galactose, starch and sucrose, fructose and mannose, and other polysaccharide metabolisms. Intriguingly, we detected the number of glycometabolism-associated pathways in host midgut at 10 dpi (4) was more than that in midgut at 7 dpi (2), suggestive of the participation of DElncRNAs in glycometabolism in host responses to *N. ceranae*. This result implied that with the stress time prolonged, the *A. m. ligustica* worker needed to transform various sugars in food into ATP as much as possible to offset the energy stolen by *N. ceranae*. Previous studies showed *N. ceranae* could inhibit honeybee cell apoptosis to give itself enough time for proliferation [85,86]. However, it is still unknown whether honeybee lncRNAs participate in regulation cell apoptosis and honeybee-*N. ceranae* interaction. In our study, we observed the up-regulation of two apoptosis inhibitor-associated genes (XM_016915940.1 and XM_006570714.2) located at up- and down-stream of DElncRNAs in, which indicated *N. ceranae* inside the host cells might adopt a lncRNA-mediated strategy to affect host cell apoptosis. But the underlying mechanism is unknown here and needs further effort. When cuticles and peritrophic membranes is breached, the pathogenic microorganism encounters a set of efficient cellular and humoral defenses including encapsulation, melanization, phagocytosis, enzymatic degration of pathogens as well as secretion of antimicrobial peptides [87,88]. In honeybees, phagocytosis and encapsulation are the two common defense mechanisms against fungi invasion [87,88]. In this current work, four, two, two, and two source genes of DElncRNA were observed to be enriched in ubiquitin-mediated proteolysis, endocytosis, lysosome,, and metabolism of xenobiotics by cytochrome P450. This result suggested these cellular immune pathways may be regulated by DElncRNAs during host *N. ceranae*-response process. Taken together, these results demonstrated that the corresponding DElncRNAs were likely to be specific regulators during *A. m. ligustica* responses to *N. ceranae* stress, and these DElncRNAs may participated the *N. ceranae*-response via interactions with their source genes.

In contrast with small ncRNAs, our understanding of the functions and regulatory mechanismsof lncRNAs is rather limited. Another manner for lncRNAs to exert their regulatory functions is to produce or interact with small RNAs [89,90]. In this work, 27 lncRNAs were detected to contain eight known miRNA precursors; additionally, 513 lncRNAs harboring 2257 novel miRNA precursors were discovered. The results indicate some *A. m. lisgustica* lncRNAs could be processed into miRNAs to exert their functions. We inferred that lncRNAs might be an important resource for identifying novel miRNAs. Some lncRNAs containing MREs have been proved to communicate with and regulate corresponding miRNA target genes via specifically competing for shared miRNAs [91,92]. Investigation of well-established miRNAs may help understand the functions of associated lncRNAs. In the present study, we detected in Am7CK vs Am7T and Am10CK vs Am10T. 106 and 143 DElncRNAs respectively interact with 76 and 98 miRNAs. Within the complex lncRNA-miRNA interaction networks, some DElncRNAs can link to the same miRNA, while some DElncRNAs could be targeted by several miRNAs (Figure 3). Several miRNAs were deeply studied such as miR-25 [93] and miR-30 [94]. Based on miR-25 mimic and inhibitor methods, Hua et al. found that miR-25 can reduce the expression of MALAT1 (metastasis associated with lung adenocarcinoma transcript 1) as a tumor suppressor in nasopharyngeal carcinoma [93]. As shown in the study conducted by Xie et al., up-regulating the expression level of miR-30a could reduce CD73’s (ecto-5’-nucleotidase) expression, thereby restraining the proliferation ability of CRC (colorectal cancer) cells and promoting the cell apoptosis as a tumor suppressor [94]. In this work, it’s detected that miR-25-x (homologous to miR-25) was the target of 11 DElncRNAs, while miR-30-x and miR-30-y (homologous to miR-30a) were targeted by 42 and 15 DElncRNAs. We inferred that these *A. m. ligustica* DElncRNAs might suppress N. ceranae via crosstalk with miR-25-x, miR-30-x, and miR-30-y. However, more experimental evidences are required.

DElncRNA-miRNA-mRNA regulation networks were further constructed and analyzed to explore the potential roles of the dysregulated lncRNAs. In this study, 59 up-regulated and 47 down-regulated lncRNAs in Am7CK vs Am7T, , and 55 up-regulated and 88 down-regulated lncRNAs in Am10CK vs Am10T were involved in regulation networks. Based on GO classifications, we detected that 54 and 95 target genes of DElncRNAs in midguts at 7 dpi and 10 dpi were both engaged in binding-associated activities, similar to the finding mentioned above. In addition, we observed that 12 and one target genes of DElncRNAs in Am7CK vs Am7T were involved in response to stimulus and cell killing, while 18 targets of DElncRNAs in Am10CK vs Am10T were enriched in response to stimulus, indicative of the involvement of corresponding DElncRNAs in host defense against *N. ceranae*. Moreover, pathway analyses demonstrated that a variety of target genes of DElncRNAs in both comparison groups were enriched in material metabolism-associated pathways, such as carbohydrate metabolism (eg. citrate cycle and galactose metabolism), lipid metabolism (eg. glycerolipid metabolism and glycerophospholipid metabolism), nucleotide metabolism (eg. pyrimidine metabolism and purine metabolism). Intriguingly, only one target (XM_001123191.4) of 20 DElncRNAs including XR_001702820.1 and XR_001703136.1 in midguts 7 dpi was detected to be associated with oxidative phosphorylation, an important way of energy metabolism; however, there was no enriched target gene of DElncRNAs in midguts 10 dpi. This indicates the participation of the aforementioned 20 DElncRNAs in regulating host energy metabolism at the early stage of *N. ceranae* stress. Additionally, target genes of DElncRNA-targeted miRNAs in Am7CK vs Am7T were engaged in three cellular immune pathways such as endocytosis (5), phagosome (1), and ubiquitin mediated proteolysis (1); and one humoral immune pathway (MAPK signaling pathway, 1). Interestingly, these four immune pathways were also enriched by target genes of DElncRNA-targeted miRNAs in Am10CK vs Am10T, with the same number. Collectively, these results demonstrated that corresponding DElncRNAs may play specific roles in regulation of the above-mentioned cellular and humoral immune pathways. We speculated that DElncRNAs can regulate the expression of target genes mediated by interactional miRNAs during the *N. ceranae*-response of *A. m.ligustica*.

## 5. Conclusions

In a nutshell, we identified 4749 known lncRNAs and 1604 novel lncRNAs in the midguts of *A. m. ligustica* workers, and showed that 111 and 146 lncRNAs were *N. ceranae*-responsive in midguts at 7 dpi and 10 dpi, respectively. These results suggest that the expression of host lncRNAs was significantly altered by the *N. ceranae* stress; part of DElncRNA were likely to participate in *N. ceranae*-response processe by regulating gene expression in *cis* or serving as miRNA precursors or ceRNAs, thus may be potential new therapeutic targets for microsporidiosis. Our data provide a rich genetic resource for further investigation of the functional roles of lncRNAs involved in *A. m. ligustica* response to *N. ceranae* infection, but also a foundation for revealing the underlying molecular mechanisms. Furthermore, this work offers a novel insight into understanding host-pathogen interaction during microsporidiosis of honeybee.

---

**Figure S1.** Experimental inoculation of *A. m. ligustica* worker with *N. ceranae*. **(A)** Microscopic observation of purified spores of *N. ceranae* (400×). **(B)** Artificial inoculation of *A. m. ligustica* worker with *N. ceranae* spores.

**Figure S2.** Bioinformatic pipeline for prediction of *A. m. ligustica* lncRNAs.

**Figure S3.** Pearson correlations between different biological repeats within every control and treatment groups.

**Figure S4.** Secondary structure of TCONS_00019779 that harbors ame-mir-927a. TCONS_00019779 is the precursor of ame-mir-927a.

**Table S1** Primers for RT-qPCR validation performed in this study.

**Table S2** Detailed information of DElncRNAs in Am7CK vs Am7T comparison group.

**Table S3** Detailed information of DElncRNAs in Am10CK vs Am10T comparison group.

**Table S4** Top 15 GO categories enriched by *cis*-regulatory target genes of DElncRNAs in Am7CK vs Am7T.

**Table S5** Top 15 GO categories enriched by *cis*-regulatory target genes of DElncRNAs in Am10CK vs Am10T.

**Table S6** Top 15 pathways enriched by *cis*-regulatory target genes of DElncRNAs in Am7CK vs Am7T.

**Table S7** Top 15 pathways enriched by *cis*-regulatory target genes of DElncRNAs in Am10CK vs Am10T.

**Table S8** Detailed information of 27 *A. m. ligustica* lncRNAs harboring 8 complete known miRNA precursors.

**Table S9** Detailed information of 513 *A. m. ligustica* lncRNAs harboring 2257 novel miRNA precursors.

**Table S10** Pairs of DElncRNAs and their target miRNAs in Am7CK vs Am7T.

**Table S11** Pairs of DElncRNAs and their target miRNAs in Am10CK vs Am10T.

## Authors’ contributions

DFC and RG designed this study. RG, HZ-C, YD, DDZ, SHG, HPW, ZWZ, CLX, and YZZ performed bioinformatic analysis and molecular experiment. DFC and RG supervised the work and contributed to preparation of the manuscript.

## Funding

This study was funded by the Earmarked Fund for Modern Agro-industry Technology Research System (CARS-44-KXJ7), the Science and Technology Planning Project of Fujian Province (2018J05042), the Education and Scientific Research Program Fujian Ministry of Education for Young Teachers (JAT170158), the Outstanding Scientific Research Manpower Fund of Fujian Agriculture and Forestry University (xjq201814), the Science and Technology Innovation Fund of Fujian Agriculture and Forestry University (CXZX2017342, CXZX2017343).

## Acknowledgements

We thank all editors and reviewers for their helpful and constructive comments.

## Conflict of interest

The authors declare that they have no conflict of interest.

## Ethical approval

This article was conducted without the use of human or animal participants.

